# Peroxidasin is associated with a mesenchymal-like transcriptional phenotype and promotes invasion in metastatic melanoma

**DOI:** 10.1101/2024.04.05.588346

**Authors:** Carlos C Smith-Díaz, Abhishek Kumar, Andrew Das, Paul Pace, Kenny Chitcholtan, Nicholas J Magon, Sultana Hossain Mehbuba Hossain, Michael R Eccles, Christine C Winterbourn, Martina Paumann-Page

## Abstract

Cutaneous melanoma is a highly invasive, heterogeneous and treatment resistant cancer. It’s ability to dynamically shift between transcriptional states or phenotypes results in an adaptive cell plasticity that may drive cancer cell invasion or the development of therapy resistance. The expression of peroxidasin (PXDN), an extracellular matrix peroxidase, has been proposed to be associated with the invasive metastatic melanoma phenotype. We have confirmed this association by analysing the transcriptomes of 70 metastatic melanoma cell lines with variable levels of PXDN expression. This analysis highlighted a strong association between high PXDN expression and the undifferentiated invasive melanoma phenotype. To assess the functional role of PXDN in melanoma invasion, we performed a knockout of PXDN in a highly invasive cell line (NZM40). PXDN knockout decreased the invasive potential by ∼50% and decreased the expression of epithelial-mesenchymal transition and invasive marker genes as determined by RNAseq and substantiated by proteomics analysis. Bioinformatics analysis of differentially expressed genes following PXDN knockout highlighted decreases in genes linked to extracellular matrix formation, organisation and degradation as well as signalling pathways such as the WNT pathway. This study provides compelling evidence that PXDN plays a functional role in melanoma invasion by promoting an invasive, mesenchymal-like transcriptional phenotype.

**Research Highlights:** PXDN expression is strongly associated with the invasive melanoma phenotype. Knockout of PXDN decreased invasion and expression of EMT marker genes concomitant with vast transcriptional changes relevant to many aspects of melanoma biology.

## Introduction

Cutaneous metastatic melanoma is a highly aggressive skin cancer that arises from the uncontrolled growth of melanocytes and causes ∼75% of skin cancer deaths (1). Early detection is crucial and surgical excision is mostly curative. However, once metastasis occurs, melanoma becomes notoriously difficult to treat. Its high invasiveness is a key characteristic which is fuelled by a remarkable plasticity that allows melanoma cells to transition between distinct differentiation states, such as the proliferative/melanocytic or invasive/mesenchymal-like/undifferentiated phenotypes (2, 3). Despite not being an epithelial cancer, the invasive/mesenchymal-like melanoma phenotype displays a striking similarity to cells having undergone epithelial-mesenchymal transition (EMT) (4–6). Melanoma phenotype plasticity enables high adaptability and heterogeneity, which contributes to metastasis, evasion of the immune system, as well as the development of therapy resistance (7, 8).

Peroxidasin (PXDN), an evolutionarily conserved, extracellular multidomain heme peroxidase enzyme (9), has been identified to be upregulated in metastatic melanoma (10). Critically, PXDN gene expression is upregulated in melanoma cell lines of a mesenchymal-like phenotype and PXDN gene silencing has been observed to reduce invasiveness, suggesting a functional role in invasion (10). Using cell lines from the New Zealand metastatic melanoma cell line panel (NZM) (11, 12), we have previously confirmed that PXDN gene and protein expression is correlated with invasive propensity (13, 14) . Our work furthermore showed that PXDN is active and continuously secreted to the extracellular space where the majority of PXDN is located.

In healthy tissues, PXDN is secreted to the basement membrane (15, 16) where it uses hydrogen peroxide to oxidise bromide to hypobromous acid (HOBr) to generate a specific covalent cross-link in the non-collagenous domain (NCD) of collagen IV (17, 18). This sulfilimine bond contributes to basement membrane stability and alters its biochemical and biophysical properties (19–22). To date, this is PXDN’s only confirmed physiological role. However, HOBr and its secondary oxidation products like bromamines are reactive oxidants that modify a multitude of biomolecules and are likely contributors to disease pathology (23–27). Detection of bromotyrosine, a stable product of the reaction of HOBr with tyrosine, shows that HOBr generated by PXDN leads to the oxidation of ECM proteins and to a lesser extent intracellular proteins (26, 28–30).

The ECM constitutes a substantial part of solid tumours and its biochemical and biophysical properties affect cell behaviour and influence proliferation, adhesion, migration and invasion. Accordingly, PXDN may contribute to invasion by modifying and remodelling the tumour ECM, thereby promoting a permissive environment for cancer cell invasion and metastasis. In addition to metastatic melanoma, PXDN is thought to contribute to the progression of numerous other types of solid cancers (31–35). However, the mechanisms underlying the role of PXDN in cancer are currently not well understood.

To further elucidate the role of PXDN in metastatic melanoma we have firstly analysed RNAseq data from NZM metastatic melanoma cell lines grouped by PXDN expression status. By comparing high PXDN expressing NZM cell lines with low or non-expressing cell lines we revealed compelling evidence that high PXDN is associated with the invasive phenotype in metastatic melanoma. To establish whether this could be of functional significance, we generated a CRISPR-Cas9 *PXDN* knockout (KO) of the NZM cell line NZM40. This is a cell line that is both highly invasive and exhibits high PXDN expression. We studied the effect of *PXDN* KO on invasion as well as on gene and protein expression using RNAseq and proteomics, respectively. We observed a reduction in invasiveness and a shift in gene expression towards a less invasive transcriptional phenotype. Overall, our data provides evidence that PXDN plays a functional role in melanoma invasion.

## Materials and methods

### New Zealand Melanoma (NZM) cell lines

NZM cell lines were originally isolated from surgical samples of human metastatic melanoma tumours (11). The cell lines were authenticated and generously provided by B. Baguley from the University of Auckland, Auckland, New Zealand. Cells were grown at 5 % oxygen concentration as described previously (13) and were tested routinely for mycoplasma (e-Myco™ VALiD Mycoplasma PCR Detection Kit). The NZM cell line panel comprises 102 cell lines with identified driver mutation status, RNAseq data of 72 cell lines (GEO submission in progress) and whole-exome sequencing data of 52 cell lines (12). We utilised RNAseq data from 70 cell lines for our analysis omitting two cell lines that had developed drug resistance.

### CRISPR-Cas9 NZM40 *PXDN* knockout generation and validation

NZM40 wild type cells (WT) were authenticated by short tandem repeat (STR) profiling. The KO cell line was generated by transfection of NZM40 with a plasmid containing Cas9 and sgRNA targeting exon 10 of PXDN (details in the supplementary section). Clone 10.34 was selected and PXDN protein expression was investigated with an in-house ELISA as described previously using an in-house developed anti-PXDN C-terminal polyclonal rabbit antiserum (13) and by western blotting using a polyclonal anti-PXDN antibody (Abbexa, abx101905) (1:1000 in 5 % bovine serum albumin) and a secondary goat anti-rabbit peroxidase antibody (1:10,000) and enhanced chemiluminescence for visualisation. A Direct Detect® infrared spectrometer was used to determine protein concentrations of cell lysates for equal protein loading of WT and KO in both the ELISA (250 μg of cell lysate protein in a volume of 50 μL and 50 μL of cell culture medium) and western blot assays (90 μg of cell lysate protein and 15 μL of medium loaded). PXDN was resolved on a 4-15% SDS-PAGE under non-reducing conditions and β-actin was used as a loading control. An antibody validation of the antibody and antiserum used is depicted in the supplementary section (Figure S4). PXDN activity was measured using stable isotope dilution liquid chromatography tandem mass spectrometry (LC-MS/MS) monitoring 3-bromotyrosine as a specific marker for HOBr (see supplementary data for details).

### Proliferation assay of NZM40 *PXDN* KO and WT

Growth characteristics of NZM40 *PXDN* KO cells were compared to WT cells using the ImageXpress PICO automated cell imaging system (Molecular Devices, LLC., CA 95134, USA). 2,500 cells were seeded in 96 well plates and proliferation was monitored by imaging cells at 24, 48, 72 and 96 hours. Fluorescent dyes Hoechst (Invitrogen™, H1399) and Sytox™ Green (Invitrogen™, S7020) were added 30 minutes before imaging (4x magnification using DAPI and FITC channels) at final concentrations of 500 µg/mL and 1 µM, respectively. Hoechst binds to the nuclei of live and dead cells whereas Sytox™ Green binds to the nuclei of dead cells only, allowing the determination of viable cells. A control was imaged 2 hours after seeding to confirm seeded cell numbers. CellReportXpress apoptosis analysis software was used to determine the number of live and dead cells. Viable cells were normalised to the seeding control (100 %) and dead cells were expressed as a percentage of the total cell number.

### Oris migration assay of NZM40 *PXDN* KO and WT

The Oris^TM^ cell migration assay was used to measure the migration of NZM40 WT and *PXDN* KO cells. 20,000 cells were seeded in 100 μL of medium with Oris stoppers in place. Cells were left to settle over night before stoppers were removed and the medium was exchanged. Migration of cells was stopped after 16 hours by fixing the cells with ice cold methanol (15 minutes) followed by three PBS washing steps and addition of DAPI (1 μg/mL) 10 minutes before the imaging of wells with the ImageXpress PICO automated cell imaging system. ImageJ was used to measure the distance migrated to calculate the migration area.

### Matrigel invasion assay of NZM40 *PXDN* KO and WT

Geltrex^TM^ LDEV-free reduced growth factor basement membrane matrix (Thermo Fisher Scientific A1413201) was defrosted on ice and diluted to 1 mg/mL in ice cold 0.01 M Tris (pH 8.0), 0.7% NaCl (150 μL) and was then added to Transwell permeable supports (Costar 3422) using precooled pipet tips. After incubation for 2 hours at 37°C to solidify the basement membrane matrix, the residual buffer was removed and 25,000 cells were seeded in 300 μL serum-free medium. 750 μL serum-containing medium was added to the bottom well and the plates were incubated for 24 hours at 37°C, 5 % CO_2_ and 5 % oxygen. Cells in the insert that did not invade were removed by gently wiping the insert with a cotton bud. Invaded cells that had attached to the bottom of the insert were fixed with ice cold methanol for 15 minutes, and were then washed and stained with DAPI for 15 minutes in a Nunc 24 well plate before imaging with the ImageXpress PICO automated cell imaging system. CellReportXpress single nuclei stain analysis software was used to determine the number of invaded cells.

### Spheroid invasion assay of NZM40 *PXDN* KO and WT

24 well plates were coated with 400 μL of 2 % agarose in PBS and left to set for 4 hours. 40,000 cells were seeded in 0.5 mL medium. Spheroids were left to form for 6 days with the medium exchanged every 2 days. 100 μL of Geltrex^TM^ LDEV-free reduced growth factor basement membrane matrix (5 mg/mL in serum-free medium) was cast on a coverslip and left to solidify for 30 minutes at 37°C. Spheroids of similar size were transferred into an Eppendorf tube and 50 μL of Geltrex (5 mg/mL in serum free medium) was added. Spheroids were pipetted onto coated coverslips and covered with residual basement membrane material. The coverslip was moved into a 6 well plate and covered with 1.5 mL of serum-containing medium. Images were taken at 24 and 48 hours and the distance invaded by the cells was measured using ImageJ by taking multiple measurements for each spheroid at each timepoint to calculate the mean invasion of three biological replicates.

### Culture conditions for RNAseq and proteomics analysis of NZM40 *PXDN* KO and WT

***RNAseq:*** NZM40 *PXDN* KO and WT cells, were grown as previously described (13), in three parallel flasks to seed three biological replicates on three consecutive days from individual cell culture flasks. The fourth and fifth biological replicates were seeded the following week on two consecutive days from two individual flasks. 700,000 cells were seeded in 10 mL of MEM-alpha medium with reduced FBS (0.75%) in a 10 cm cell culture dish for 48 hours. On average, the cells had proliferated to approximately 1.1 million cells at this point. After 48 hours, cells were replenished with fresh 0.75% FBS medium for 5 hours before the medium was fully removed and replaced with 6 mL medium containing 0% FBS. After 22 hours, the cells were washed twice with PBS, the buffer was removed and the cells were harvested by scraping in residual PBS. The cells were snap frozen on dry ice and stored at -80 °C for a short period of time until RNA extraction.

#### Proteomics

NZM40 *PXDN* KO and WT cells were grown as for the RNAseq experiments and were harvested 24 hours after exchange to medium with 0% FBS. The medium was removed from cells and spun at 400 g to remove any detached cells. The supernatant contained secreted proteins and was snap frozen on dry ice and stored at -80 °C. The cells were washed three times in PBS and scraped in residual buffer, snap frozen and stored at -80 °C. Three biological replicates of NZM40 *PXDN* KO and WT cells were performed.

### RNA extraction and sequencing of NZM40 *PXDN* KO and WT

RNA was isolated using the NucleoSpin^®^ RNA Plus kit following the manufacturer’s protocol (Macherey-Nagel, Düren, Germany). The integrity and concentration of RNA samples was assessed using the TapeStation 2200 (Agilent, Santa Clara, California) in house. Purified RNA samples were sent to China for quality control, library preparation and sequencing (Annoroad Gene Technology Co., Ltd. Beijing, China). RNA was purified using oligo(dT) beads and library preparation was carried out with the Ultra Directional RNA Library Prep Kit for Illumina (New England Biolabs, Ipswich, MA, USA). The Illumina NovaSeq6000 platform was used for library sequencing. Each sample was sequenced at approximately 20 million reads (using 150 bp paired end reads).

### Label-free quantitative proteomics of NZM40 *PXDN* KO and WT

Proteomic analysis of cell lysate samples and cell medium samples containing the secretome was performed using a label-free quantitative approach as described in detail in the supplementary section. Briefly, protein estimation assays were used to normalize protein content between samples before cell lysates and cell medium samples were processed, reduced, alkylated and tryptically digested. Both cell and secretome samples were chromatographically separated on an emitter-tip column packed with 3 µM C-18 Luna material (Phenomenex, USA)) on an Ultimate 3000 uHPLC (Thermo Scientific, USA) coupled to the LTQ Orbitrap XL mass spectrometer (Thermo Scientific, USA). Each biological repeat was measured in 3 technical replicates. The resulting data from all samples were analysed with the Proteome Discoverer (PD) software (version: 2.5, Thermo Scientific, USA). The data was searched against the human proteome (downloaded: February 2022, Uniprot.org) with the Sequest HT search engine node inside the PD software. The search was set up to look for semi-tryptic peptides. The resulting quantitative data were normalised on the sum of abundances from all peptides detected from all samples. The relative abundance of the proteins was calculated with the Top-3 approach where the average abundance of the three most abundant peptides for a particular protein was used. The resulting abundance values were used to calculate the protein ratio between NZM40 *PXDN* KO to WT to generate a list of differentially abundant proteins.

### Bioinformatics

RNAseq FASTQ files from the NZM cell lines (12) (GEO submission in progress) were trimmed using TrimGalore! and mapped to the human genome Hg38 using Hisat2. BAM files were imported into Seqmonk (Babraham Bioinformatics, Cambridge, UK) and NZM cells lines were grouped by PXDN expression status. The 22 cell lines with highest PXDN expression (log_2_ RPM 7.64 to log_2_ RPM 11.72) were assigned to the “high PXDN group” and the 22 cell lines with lowest PXDN expression (log_2_ RPM -4.74 to log_2_ RPM 0.09) were assigned to the “low PXDN group”. Differential gene expression analysis was performed between groups in Seqmonk using the DESeq2 Bioconductor package (FDR < 0.05) with further downstream analysis in R.

RNAseq data from *PXDN* KO experiments was processed in a similar manner although adapters and low-quality reads were removed using Cutadapt (v1.9.1). Quality control was performed post-trimming using FASTQC (Babraham Bioinformatics, Cambridge), with mean per base sequence quality scores at the 150 bp position > 34 for all samples. Mapping quality control was performed in Seqmonk with >99% of reads falling within genes and >96% within exons. Transcriptomes were grouped by treatment group and differential gene expression between groups was performed using DESeq2 (FDR < 0.05). Overrepresentation analysis was performed in R using ReactomePA, and heatmaps were generated using pheatmap. Gene set enrichment analysis (GSEA) was performed in R with FGSEA using the shrunk log2 fold change (DESeq2) as the ranking metric (36, 37). For GSEA, gene sets from MSigDB Hallmark 2020 as well as manually curated melanoma phenotype datasets were used (3, 38). RNAseq data was uploaded to GEO (accession number: GSE259402).

## RESULTS

### RNAseq analysis of NZM cell lines by PXDN gene expression

Comparing the transcriptomes of 70 NZM cell lines revealed high heterogeneity in PXDN expression (Figure 1A). To determine transcriptional patterns associated with high and low PXDN expression in melanoma, cell lines with high PXDN expression (n=22) were compared against cells with low PXDN expression (n=22), omitting transcriptomes with intermediate levels of PXDN. When these 44 cell lines were compared, principal component analysis (PCA) highlighted clustering of cells by PXDN expression status along PC1, suggesting that PXDN expression is associated with different transcriptional states in NZM cells (Figure 1B). However, the clustering apparent along PC1 only represents a moderate effect given the relatively low degree of variance captured by PC1 and degree of overlapping transcriptomes. Further, differential gene expression analysis (FDR <0.05) highlighted 1909 differentially expressed genes between high and low PXDN groups, with 1370 downregulated and 539 upregulated genes in low relative to high PXDN expressing cells (Figure 1C).

**Figure 1:**
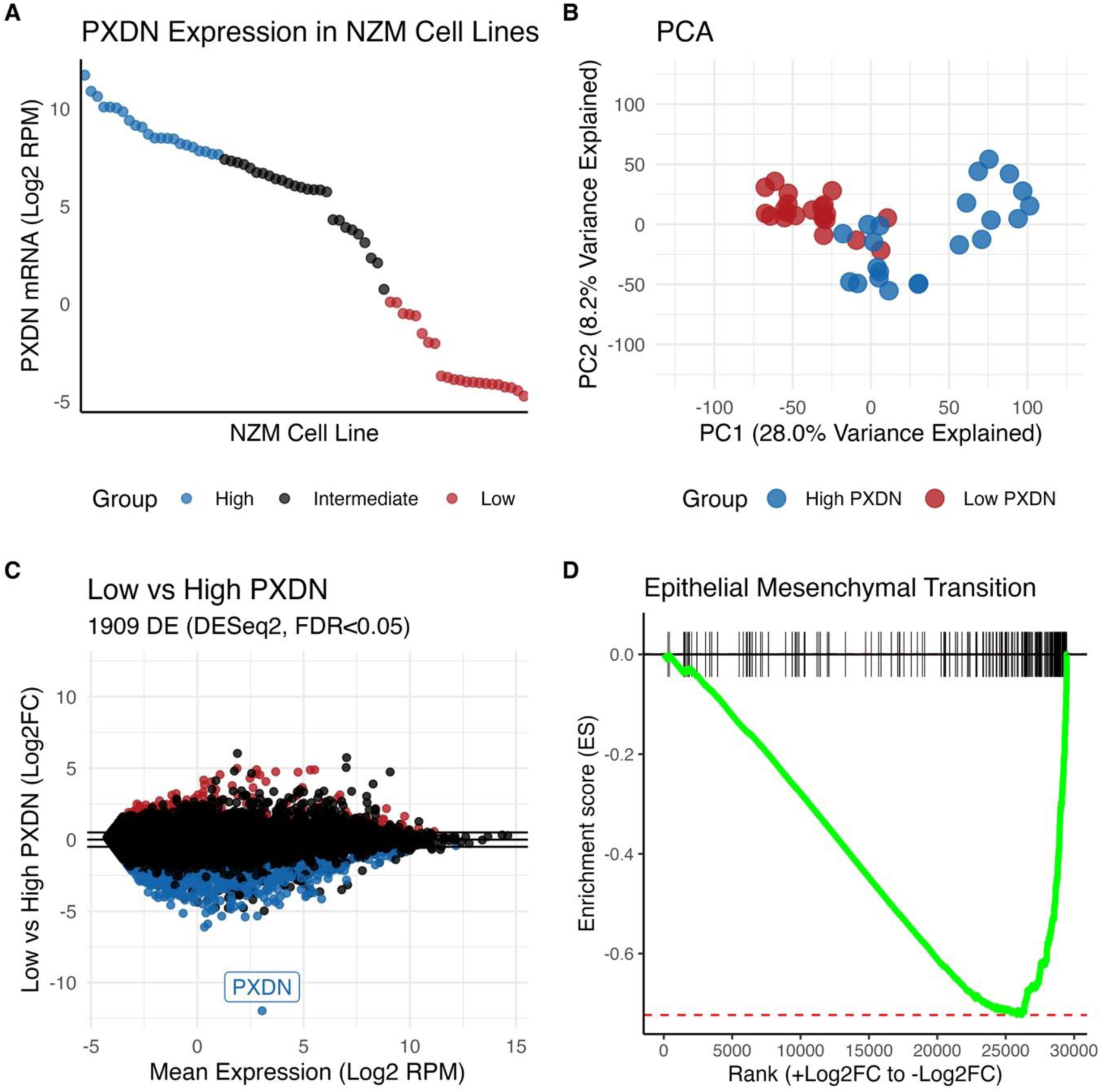
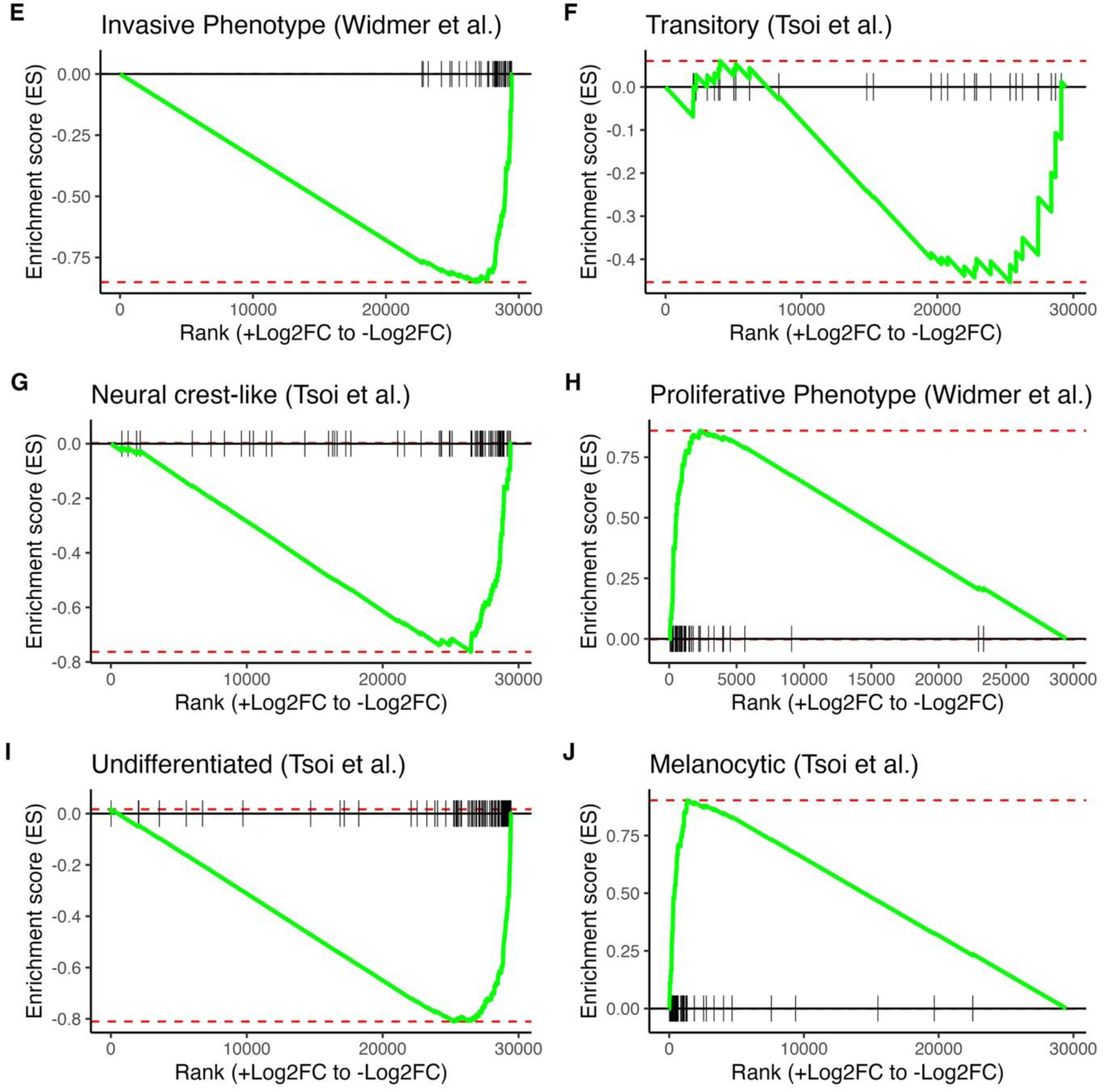
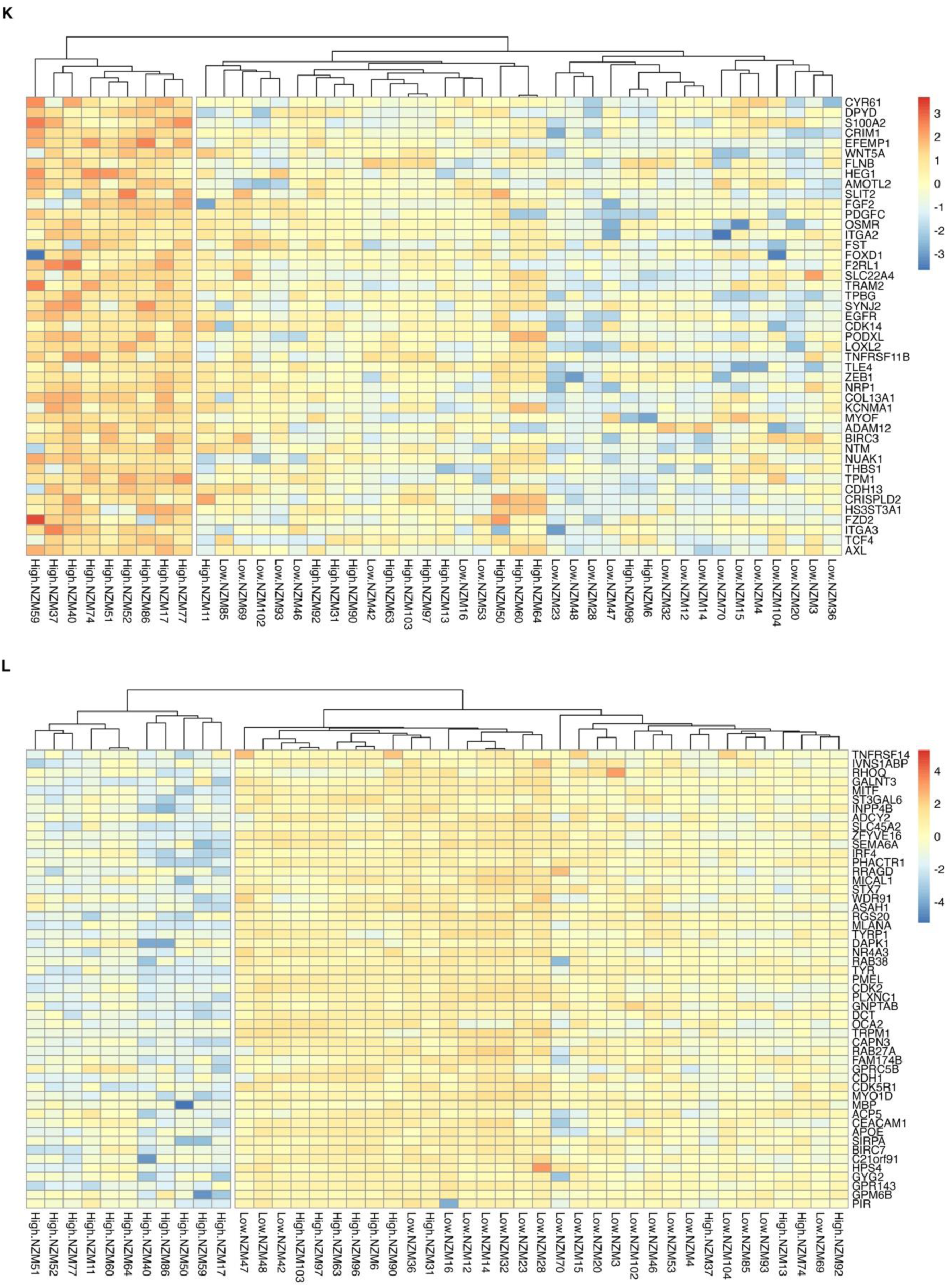
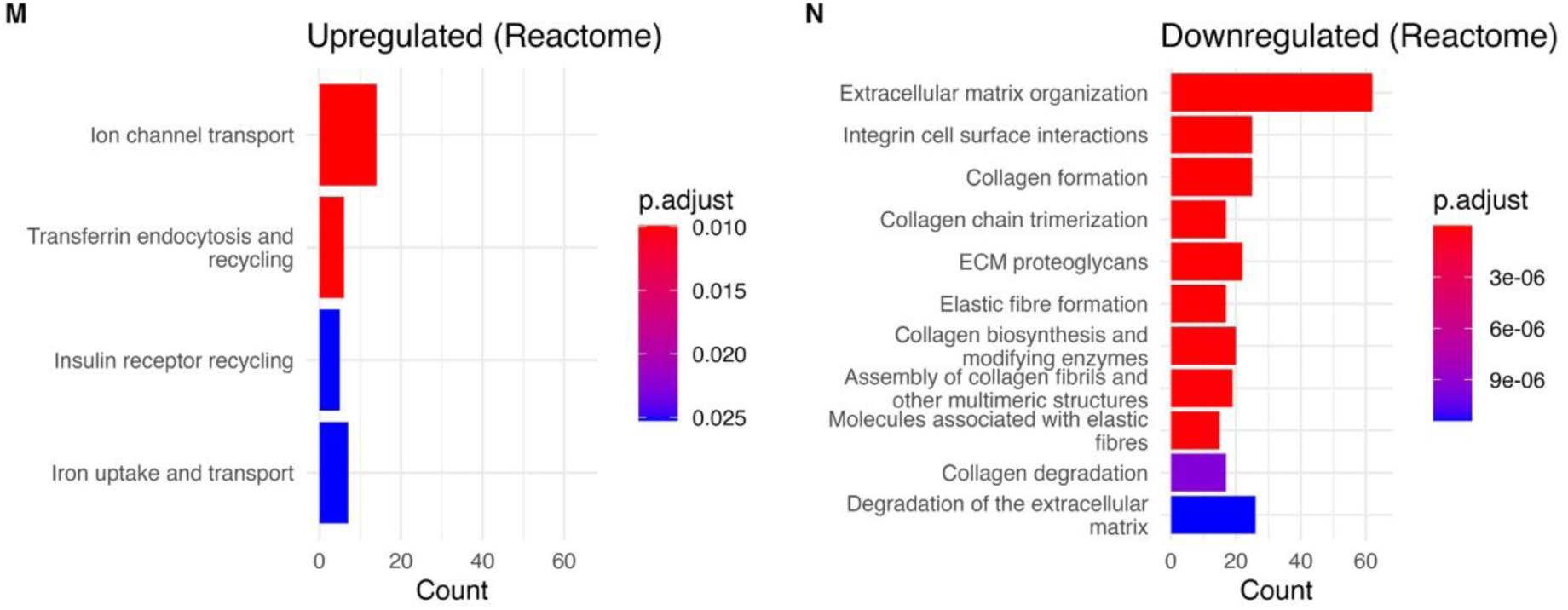
Analysis of transcriptional differences between NZM cells with low vs high PXDN expression. **A:** NZM cell lines were grouped by PXDN expression status (n=70, RPM, reads assigned per million mapped reads). The top 22 cell lines were assigned to the “High” group whereas the 22 cell lines with lowest PXDN expression were allocated to the “Low” group. **B:** PCA analysis showing clustering of the 44 selected cell lines by PXDN expression along PC1. **C:** Differential gene expression analysis (DESeq2 FDR <0.05) revealed 1909 differentially expressed genes (red = upregulated, blue = downregulated) between low and high PXDN expression groups. **D:** GSEA was performed using FGSEA, highlighting a decrease in the EMT (epithelial-mesenchymal transition, MSigDB_Hallmark_2020) pathway in low PXDN expressing cell lines (padj = 4.78 x10^-22^, enrichment score = -0.72). **E-J**: GSEA was performed using a manually curated database of melanoma phenotype gene sets identified by Widmer et al. and Tsoi et al. Comparing transcriptional differences between low and high PXDN expression groups highlighted a negative association between low PXDN associated genes and the (E) invasive (padj = 9.46 x 10^-13^), (I) undifferentiated (padj = 7.59 x 10^-25^) and (G) neural crest-like (padj = 2.20 x 10^-10^) phenotypes with no statistically significant enrichment for the transitory phenotype (F). In contrast, there was a positive association between low PXDN associated genes and the (H) proliferative (padj = 1.64 x 10^-18^) and (J) melanocytic (padj = 4.50 x 10^-28^) phenotypes. **K-L**: Heatmaps depicting the relative expression of invasion (K) and proliferation (L) hallmark genes for the 44 selected high and low PXDN expressing NZM cell lines. **M:** Pathway analysis (over-representation test, ReactomePA, R) on upregulated genes in low PXDN expressing cell lines. **N:** Pathway analysis on downregulated genes in low PXDN expressing cell lines. Files of tables depicting differentially expressed genes and pathway analysis outputs are available in the supplementary section.

To compare the association between PXDN expression and different melanoma phenotypes we compared the expression change of genes in selected melanoma pathways between low and high PXDN groups. We performed gene set enrichment analysis (GSEA) to show that genes in the epithelial-mesenchymal transition pathway (EMT, MSigDB_Hallmark_2020) (Figure 1D) and genes linked to the invasive/undifferentiated and neural crest-like melanoma phenotypes were positively correlated with high PXDN expression (Figure 1E, 1G, 1I). In contrast, genes associated with the proliferative/melanocytic non-invasive state were positively correlated with low PXDN expression (Figure 1H and 1J) (3, 38). This analysis suggested an association between increased PXDN expression and the transition from a non-invasive, melanocytic phenotype towards an invasive, undifferentiated phenotype.

We then examined the relative expression of hallmark genes representing the invasive or proliferative melanoma phenotypes (38) at the individual cell line level. This revealed clustering of individual cell lines by PXDN expression across gene sets characteristic of invasive or proliferative phenotypes (Figure 1K, 1L). The heatmaps viewed as a whole, showed clear associations between PXDN expression and different melanoma phenotypes (invasive or proliferative). For example, a cluster of cell lines with high PXDN expression exhibited very high overall expression of hallmark genes for the invasive phenotype, and an overlapping cluster of high PXDN transcriptomes exhibited very low expression of proliferative phenotype hallmark genes. Some cell lines did not follow this trend (i.e. NZM96 or NZM6, cell lines with relatively high PXDN expression but low expression of invasive hallmark genes). This indicates that PXDN is not the sole determinant of melanoma phenotype. However, it does display a strong association with the invasive transcriptional state.

Applying overrepresentation analysis using the Reactome database confirmed these melanoma phenotype associations (Figure 1M, 1N). Genes downregulated in low PXDN expressing cells were highly enriched for biological processes pertaining to extracellular matrix remodelling, as might be observed during epithelial-mesenchymal transition. Many of the top downregulated pathways related to ECM organisation, processes affecting collagen and elastic fibre formation, collagen biosynthesis and trimerisation, ECM proteoglycans and integrin cell surface interactions. In contrast, overrepresentation analysis on the upregulated genes in low PXDN expressing cells did not reveal any compelling enriched pathways. Overall, this analysis of 44 metastatic melanoma cell lines identifies PXDN as a novel biomarker of invasiveness and the undifferentiated or neural crest-like phenotypes.

### Characterisation of PXDN CRISPR-Cas9 knockout of high PXDN expressing cell line NZM40

To investigate whether PXDN expression and melanoma invasion were functionally connected, we generated a *PXDN* knockout (KO) cell line and compared its properties with wild type (WT) cells. A knockout of the high PXDN expressing cell line NZM40 was generated using CRISPR-Cas9 to edit exon 10 of PXDN, targeting the second Ig-domain (Figure S1A&B). PCR sequencing confirmed biallelic editing of the respective sequence, leading to the introduction of a stop codon (Figure S1C). Surprisingly, a low residual level of PXDN protein was still detectable in the knockout. Comparing PXDN protein expression of NZM40 *PXDN* KO with the wild type (WT) using an ELISA (Figure 2A) and western blotting (Figure 2B&C) showed a decrease in PXDN expression of about 90%. Only a small fraction of the residual protein was secreted (2A,B), whereas in the WT about 80% of PXDN was located extracellularly. Approximately 25% of intracellular PXDN (Figure 2A, black) was detectable in the KO compared to the WT. Western botting showed that intracellular PXDN was not truncated in the KO and densitometry indicated the loss of approximately 90% of the protein (Figure 2C). The residual protein was catalytically active, as shown in Figure 2D using a mass spectrometric assay detecting bromotyrosine, which is a specific marker for HOBr generated by PXDN from hydrogen peroxide and bromide. This indicates an intact active site structure of the residual protein. NZM40 *PXDN* KO did not affect rate of proliferation (Figure 2E) or the number of cells undergoing cell death (Figure 2F).

**Figure 2:**
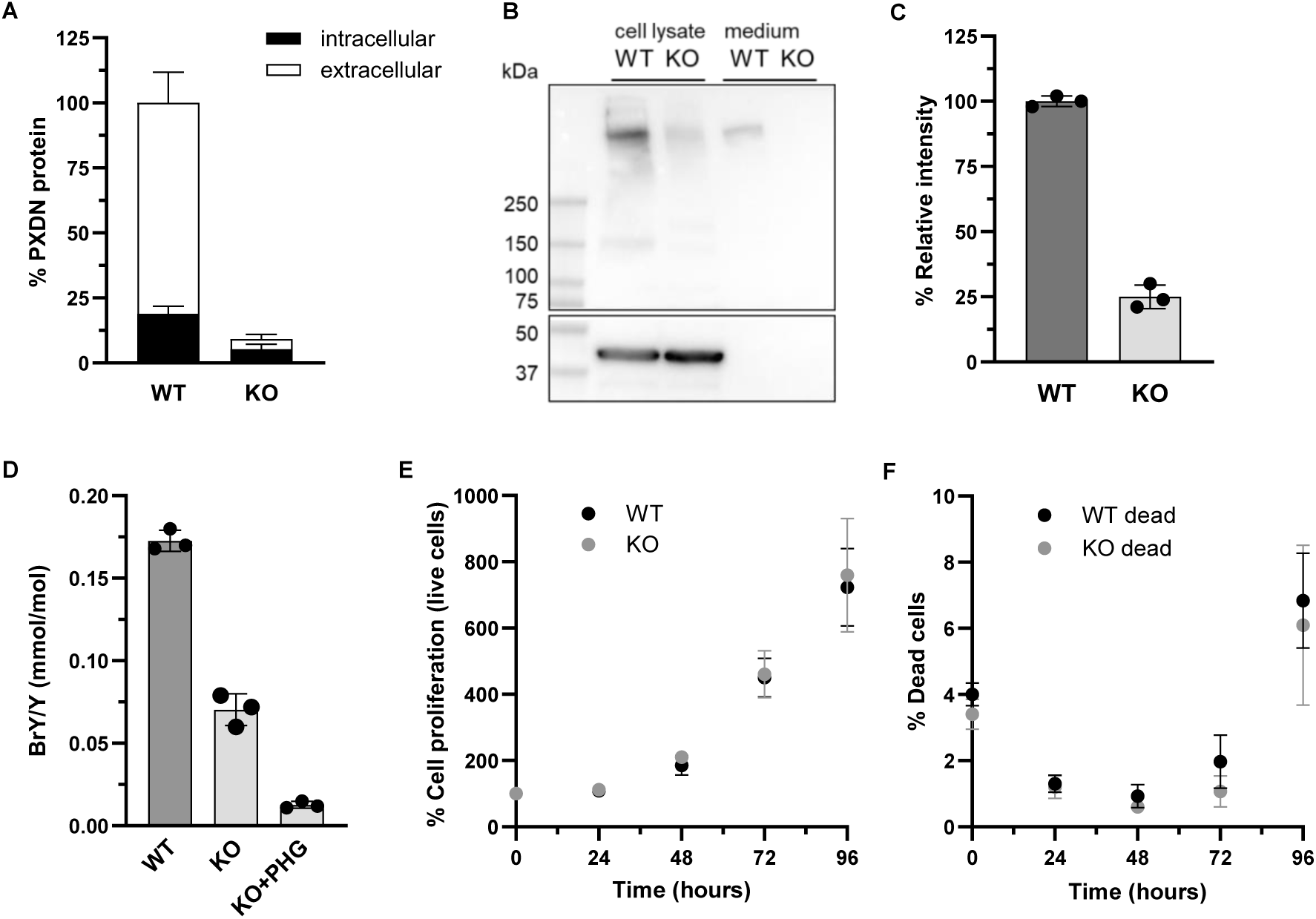
PXDN protein expression, catalytic activity and proliferation of NZM40 WT and *PXDN* KO. **A:** Extra- and intracellular expression of PXDN protein in the NZM40 WT and *PXDN* KO cell lines measured by ELISA. Bars represent mean values of three individual experiments ± SD. **B:** Representative western blot of PXDN protein expression of NZM40 WT and *PXDN* KO cell lines under non-reducing conditions depicting the ∼500 kDa homotrimer of PXDN and a faint band for the not yet trimerized monomer. Loading control β-actin is shown in the lower panel. **C:** Densitometry analysis (bars represent mean ± SD of three experiments shown by individual data points) of 500 kDa PXDN band of western blot. **D:** Bromination activity of NZM40 WT, *PXDN* KO and inhibition of *PXDN* KO activity with peroxidase inhibitor phloroglucinol (PHG). Individual data points of three experiments, bars represent the mean value ± SD. **E:** Proliferation of NZM40 WT and *PXDN* KO cell lines seeded in 96 well plates and cell numbers were determined using the Pico Imaging Express 2 hours after seeding and at 24, 48, 72 and 96 hours, counting live and **F:** % dead cells. Datapoints represent the mean ± SD of 3 independent experiments. Cell numbers were normalised to cell count at 2 hours.

### Invasion, migration and morphology of NZM40 PXDN KO compared with WT cells

Both matrigel and spheroid invasion assays showed that NZM40 *PXDN* KO cells exhibited an approximately 50% decrease in invasive potential when compared to WT cells (Figure 3A and 3B respectively). Interestingly, *PXDN* KO had no impact on the migratory capacity of the cells (Figure 3C). During early passages, and when grown to a high density, NZM40 *PXDN* KO cells displayed a cobblestone-like morphology with an increase in cell-cell interactions, which is characteristic of epithelial-like cells. In contrast, PXDN WT cells displayed more of a spindly mesenchymal-like cell morphology (Figure 3D).

**Figure 3:**
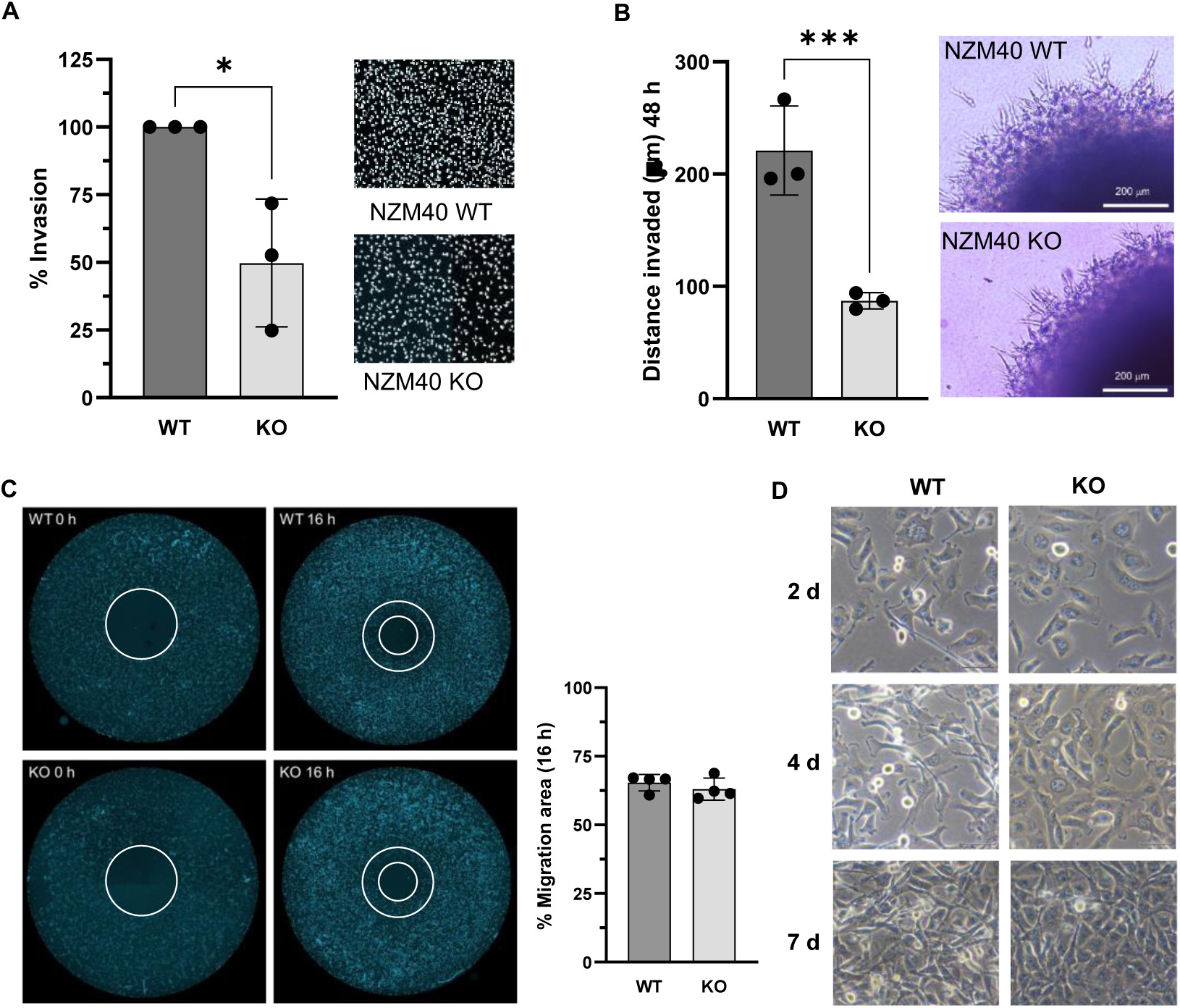
Matrigel and spheroid invasion, migration and cell morphology of NZM40 WT and *PXDN* KO. **A:** Matrigel invasion assay (1 mg/mL Matrigel) fixed and stained 24 hours after seeding of NZM40 WT and *PXDN* KO. The number of KO cells that invaded were expressed as a percentage of the number of WT cells of three biological replicates shown as individual data points. Bars represent the mean ± SD *, *P* <0.05 by unpaired, two-tailed t-test. **B:** Spheroid invasion assay depicting the invaded distance at 48 hours of NZM40 WT and *PXDN* KO. ***, *P* <0.001 by unpaired, two-tailed t-test. **C:** Representative DAPI stained migration images of NZM40 WT and *PXDN* KO at indicated times using the ORIS migration assay. The circle at time points zero depicts the cell free area immediately after the removal of the insert that blocked cell growth within the circle. 16 hours later the cells migrated into the circle and reduced the cell free area to the size of the inner circle. The cell free area was determined using ImageJ and the area the cells had migrated into was calculated based on the % relative to the cell free area at time zero. Error bars represent the mean of individual experiments depicted as single data points (n=4) ± SD. **D:** The morphology of *PXDN* KO compared to WT showed an increase in cell-cell interactions indicative of a more epithelial-like phenotype.

### RNAseq analysis of NZM40 PXDN KO gene expression compared to WT

RNAseq analysis of NZM40 *PXDN* KO vs NZM40 WT cell lines identified large transcriptional differences that were consistent between replicates. 6091 genes were differentially expressed (FDR < 0.05) (Figure 4A) with 3191 downregulated and 2900 upregulated genes in *PXDN* KO relative to WT cells. Clear clustering between KO and WT transcriptomes was apparent with PCA analysis (Figure 4B). Interestingly, despite the CRISPR-Cas9 induced mutation in the *PXDN* KO being expected to impair translation and to only manifest in altered protein abundance, total levels of PXDN mRNA were also highly downregulated in KO cells (Log2FC -3.5, Figure 4A).

**Figure 4:**
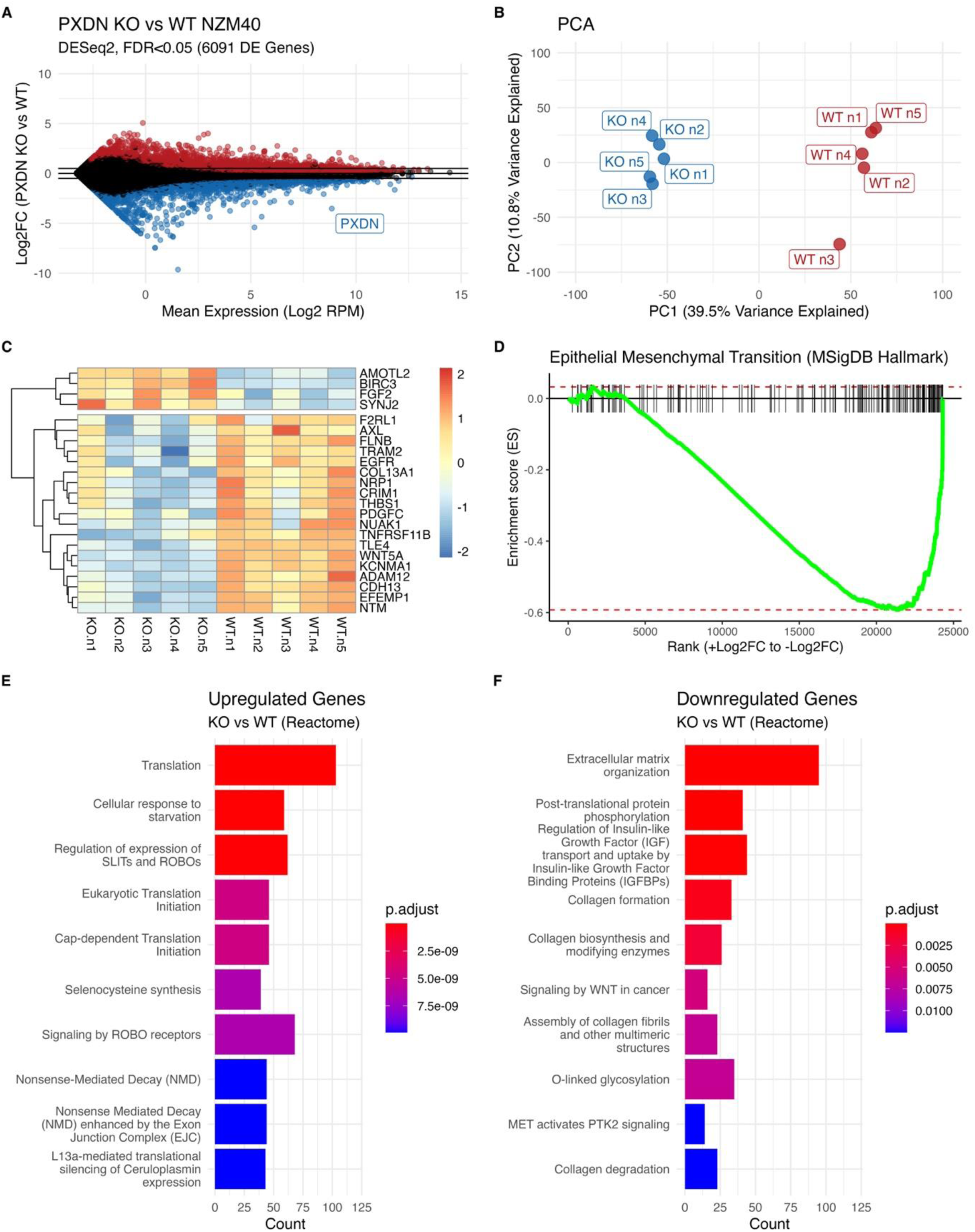
Analysis of transcriptional differences between NZM40 *PXDN* KO and WT cells. **A:** Differential gene expression analysis (DESeq2, FDR <0.05) was performed between *PXDN* KO and PXDN WT NZM40 cell lines (n=5), revealing 6091 differentially expressed genes. Genes in red represent upregulated genes, and genes in blue represent downregulated genes in the *PXDN* KO vs WT comparison. **B:** PCA analysis highlights clear clustering of five biological repeats of *PXDN* KO and WT along PC1. **C:** Genes identified as representing the invasive melanoma phenotype by Widmer *et al.* with differential expression in KO vs WT cells (DESeq2 < 0.05) depicted in heatmap format. This illustrates a net decrease in the expression of invasive phenotype genes following PXDN knockout. **D:** Gene set enrichment analysis (GSEA) was performed with FGSEA using a dataset of EMT (epithelial-mesenchymal transition) signature genes from MSigDB_Hallmark_2020. After ranking genes by expression change (shrunk log2 fold change, DESeq2) between *PXDN* KO vs WT groups, downregulated genes in the *PXDN* KO group were highly enriched for EMT signature genes (padj = 5.43 x10^-10^, enrichment score -0.58). **E:** Unbiased pathway analysis (over-representation test, ReactomePA, R) on upregulated genes in *PXDN* KO vs WT cells. **F:** Unbiased pathway analysis (over-representation test, ReactomePA, R) on downregulated genes in *PXDN* KO vs WT cells. This reveals a strong enrichment of pathways linked to the extracellular matrix and collagen remodelling. Files of tables depicting differentially expressed genes and pathway analysis outputs are available in the supplementary section.

We examined the expression of invasive phenotype hallmark genes identified by Widmer *et al.* that were also differentially expressed between KO and WT cell lines and found that the majority (19/23 genes) were downregulated in *PXDN* KO cells (Figure 4C). A similar trend was also apparent when all genes linked to the invasive phenotype were considered (irrespective of differential expression) (Supplementary Figure S2A-B). Gene set enrichment analysis (GSEA) of genes in the epithelial-mesenchymal transition pathway (EMT, MSigDB_Hallmark_2020) showed mostly downregulated gene expression in the *PXDN* KO compared to the WT (Figure 4D). The top downregulated genes by fold change following *PXDN* KO were *DKK1* (Dickkopf WNT Signalling Pathway Inhibitor 1, Log2 FC -9.34), and *FZD8* (Frizzled Class Receptor 8, Log2FC -7.46) – both genes linked to WNT signalling and EMT. Other examples of highly downregulated EMT related genes were *CXCL12* (C-X-C motif chemokine ligand 12, Log2FC -6.28), *MXRA5* (matrix remodeling associated 5, Log2FC -6.00), *ZEB2* (zinc finger E-box binding homeobox 2, Log2FC -5.70), *THBS2* (thrombospondin 2, Log2FC -5.41) and *LUM* (Lumican, Log2FC 4.57). E-cadherin (*CDH1* Log2FC 2.57) was upregulated in the KO whereas N-Cadherin was downregulated (*CDH2* Log2FC -1.28).

We then performed overrepresentation analysis of differentially expressed genes using the Reactome database and observed that downregulated genes were enriched for processes pertaining to the extracellular matrix, collagen formation/degradation and WNT signalling in cancer. Crucially, the enriched downregulated pathways in KO cells pertaining to the ECM closely mirrored those enriched in downregulated genes in low PXDN expressing NZM cell lines (Figure 1E&F, Supplementary Figure S3A-B and the supplementary file depicting the overlap of genes in PXDN KO and low PXDN expressing cell lines). Conversely, many of the enriched upregulated pathways in *PXDN* KO cells were involved in the regulation of translation (Figures 4E-F).

### Proteomic analysis of NZM40 PXDN KO protein expression compared to WT

To complement the RNAseq data, we examined changes in protein abundance in the cell lysate (Figure 5A) and secretome (Figure 5B) arising from *PXDN* KO. We identified 48 proteins with differential abundance (log2 FC > |1|, p <0.05) in the secretome. Only 13 differentially expressed proteins (log2 FC > |1|, p <0.05) were identified in the lysate. Consistent with the specific protein analyses in Figure 2, PXDN was highly downregulated in the secretome, and to a lesser extent in the lysate. We performed overrepresentation analysis on the upregulated and downregulated pathways in the secretome following *PXDN* KO and again observed downregulated pathways pertaining to extracellular matrix organisation, degradation and collagen structures (Figures 5C-D). Considering all the proteins identified as having differential abundance (p<0.05) in KO vs WT cells, there was a strong correlation with mRNA expression changes in RNAseq datasets (Figure 5E).

**Figure 5:**
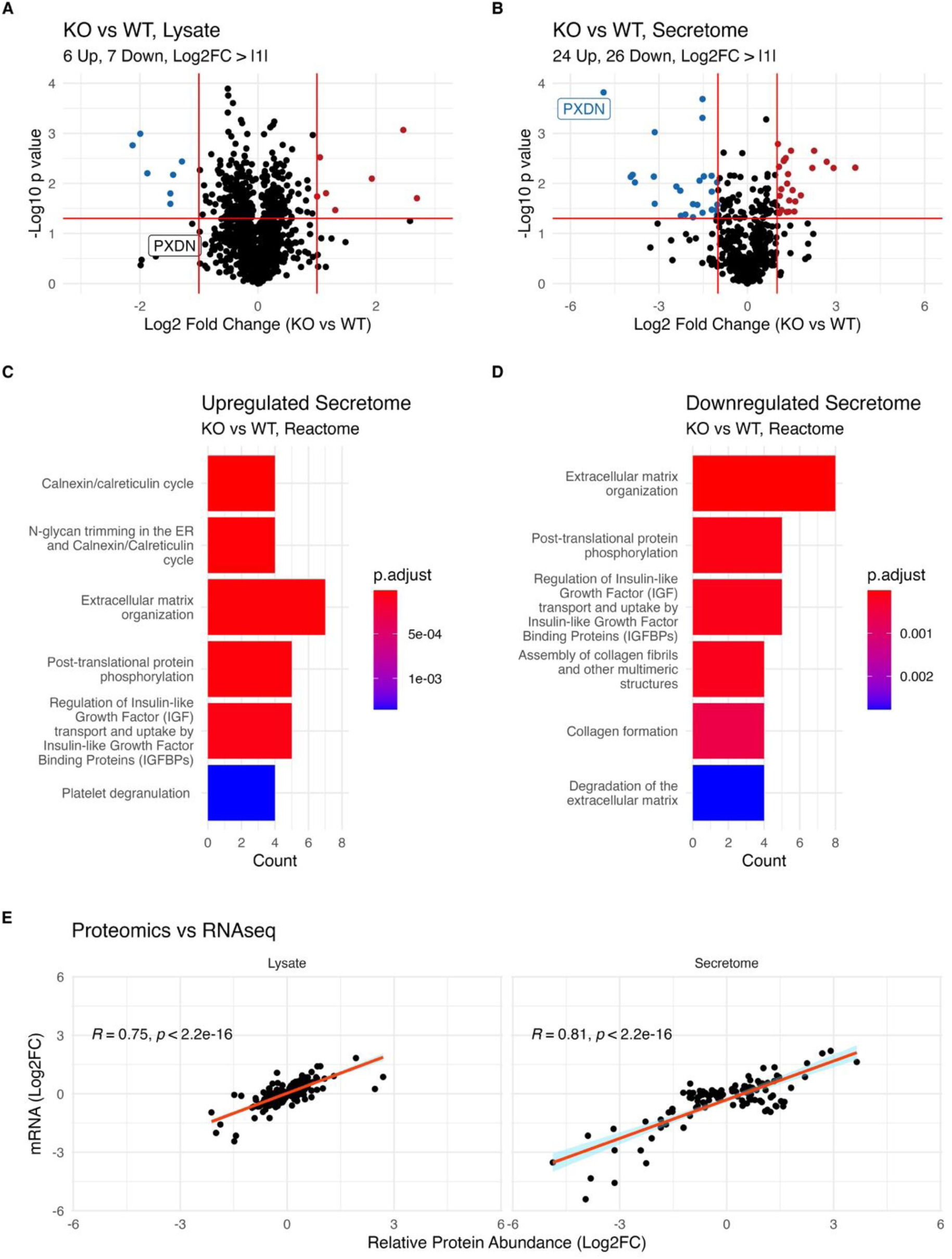
Proteomics analysis of secretome and lysate of *PXDN* KO vs WT. **A:** Differentially expressed proteins in the secretome and **B:** lysate of *PXDN* KO (downregulated proteins in blue, upregulated in red; log2 FC > |1|, p <0.05). Overall, 1525 proteins were detected in the lysate and 681 were detected in the secretome. **C-D:** Unbiased pathway analysis (over-representation test, ReactomePA, R) of upregulated and downregulated proteins in *PXDN* KO vs WT cells. **E:** Correlation between differentially abundant proteins from proteomic experiments (p < 0.05) and the gene expression change of corresponding genes in the transcriptomic datasets shown at Figure 4. Files of tables depicting differentially expressed proteins and pathway analysis outputs are available in the supplementary data.

### Association between PXDN and melanoma transcriptional phenotypes

To further explore the associations between PXDN expression and different transcriptional phenotypes in melanoma, we selected the entire panel of NZM transcriptomics data and the RNAseq data from the experiments we conducted with NZM40 and NZM40 PXDN KO cells. Transcriptomics data from the two datasets were processed separately, PXDN expression was quantified as RPKM (without any log2 transformation) and the mean expression values (RPKM) for gene sets representing the invasive, proliferative, melanocytic and undifferentiated transcriptional phenotypes were calculated. We then compared PXDN expression with different melanoma phenotypes across these different transcriptomes. As we would expect, these datasets show that PXDN expression is positively correlated with the invasive and undifferentiated phenotypes and negatively associated with the proliferative and melanocytic phenotypes. Intriguingly, all cells with very high PXDN expression exhibit higher expression of undifferentiated phenotype genes, whereas around half the cells with high expression of invasive phenotype genes also have high PXDN expression. Notably, none of the cell lines with high expression of genes linked to proliferative or melanocytic phenotypes have high PXDN expression (**Figure 6**). This suggests that melanoma cells that originate in the proliferative/melanocytic state have low basal PXDN expression. During the process of “dedifferentiation” towards the undifferentiated state, melanoma cells may upregulate PXDN expression, which is a hallmark of some (but crucially not all) transcriptionally invasive cells.

**Figure 6:**
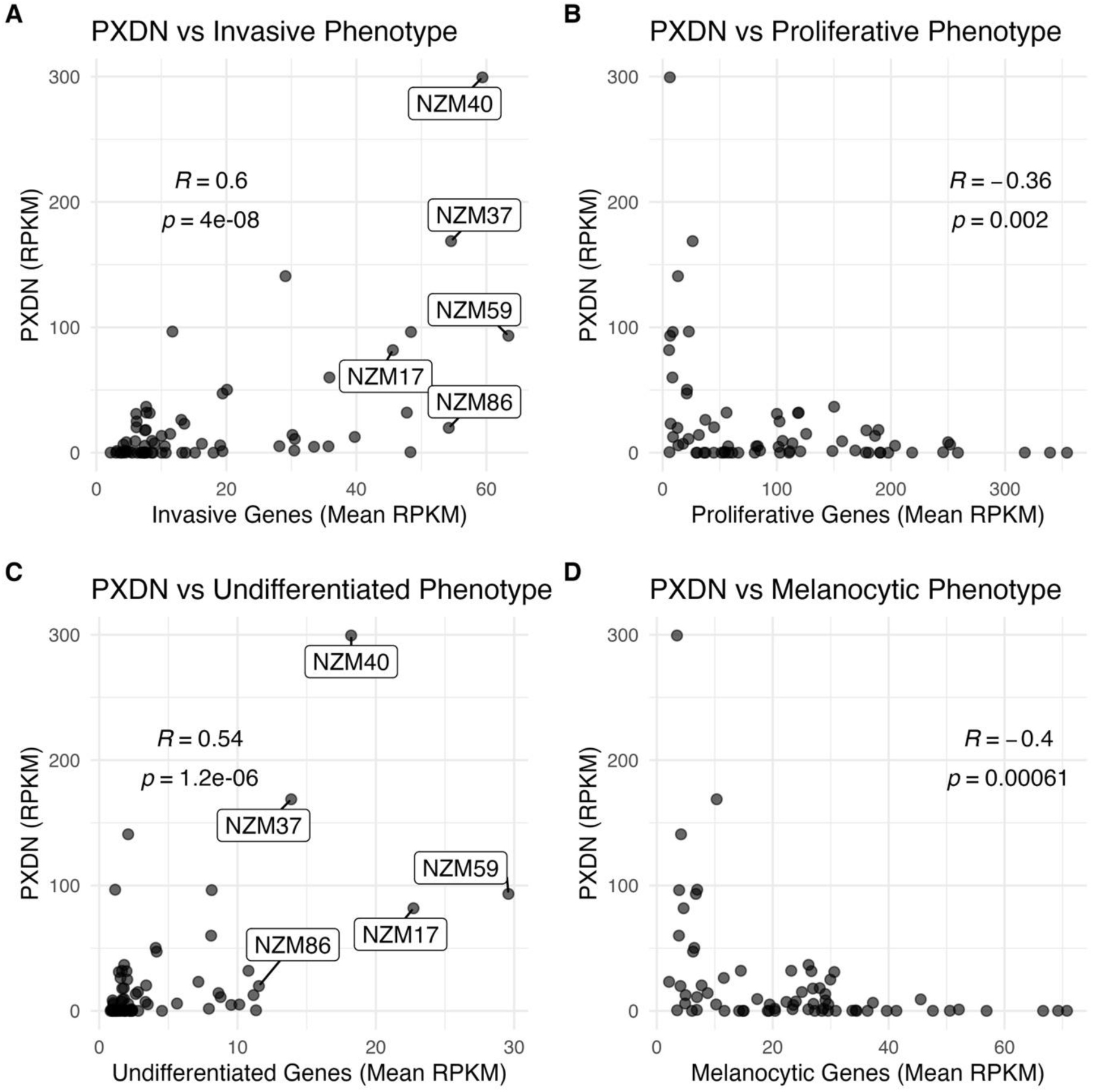
Association between PXDN expression and melanoma transcriptional phenotypes. **A-D**: The expression of PXDN in the NZM panel was compared with the mean expression of genes from the curated database of melanoma phenotype genes identified by Widmer *et al.* and Tsoi *et al.* Pearson’s correlation coefficient is report from each graph, calculated in R using stat_cor(method = "pearson").

Notably, some differences in PXDN expression (and invasive gene expression) are apparent between NZM40 transcriptomes from different sequencing experiments (i.e. the data reported at Figure 1 vs Figure 4). A variety of technical and biological effects may underpin these differences (such as different culture conditions). However, even within NZM40 cells from different experiments, including the PXDN KO cells, PXDN expression is proportional to invasive gene expression. In the NZM40 PXDN KO we see a decrease in invasive gene expression as well as an increase in undifferentiated gene expression, however, there is no reversion back towards the melanocytic / proliferative.

## DISCUSSION

The treatment of metastatic melanoma remains highly challenging due to its heterogeneity, genetic instability, high invasive and metastatic potential and its ability to switch between different phenotypes. All of these traits contribute to disease progression, the development of resistance to standard therapies and often, to disease relapse (39). Consequently, identifying and targeting the specific effectors that mediate melanoma progression is critical.

Our study provides novel insights into the role of PXDN in melanoma. Firstly, our analysis of 44 metastatic melanoma transcriptomes from high and low PXDN expressing cell lines confirmed the strong association between PXDN expression and the invasive melanoma phenotype, which we previously reported for eight NZM melanoma cell lines (13, 14). Gene set enrichment analysis using three independently identified sets of maker genes (EMT genes from MSigDB_Hallmark_2020; the invasive and proliferative phenotype marker genes identified by Widmer et al. (38); and undifferentiated, neural crest-like, transitory and melanocytic phenotype marker genes identified by Tsoi et al.(3)) provides compelling evidence that PXDN is associated with EMT and the invasive, undifferentiated and neural crest-like phenotypes. PXDN expression is not the sole determinant of cell phenotype and not all transcriptionally invasive melanoma cells have high PXDN expression. This indicates the complexity of melanoma and suggests that PXDN expression is likely one of many factors facilitating melanoma invasion. For example there are some cell lines (i.e NZM6 or NZM96) that displayed high PXDN expression but lacked invasive transcriptional characteristics. Nevertheless, we have observed that all NZM cells with a proliferative / melanocytic phenotype have low PXDN expression. Notably, despite some heterogeneity, most of the NZM cells that have transitioned to an undifferentiated state have acquired high PXDN expression. Overall, PXDN expression is strongly associated with the invasive phenotype, and the gene represents a novel biomarker of the invasive phenotype in melanoma.

As an association does not necessarily represent a causal relationship we generated a *PXDN* KO cell line (NZM40 KO). With ∼10% residual PXDN protein expression in the KO, we observed a reduction of the invasive potential by ∼50%, suggesting that PXDN has a functional role in melanoma invasion. Residual PXDN protein expression in the KO could be due to stop codon readthrough (40) or biological plasticity partially rescuing CRISPR-Cas9 knockouts (41). *PXDN* KO showed an almost complete loss of secreted PXDN as confirmed by our proteomics data. This suggests that extracellular PXDN may be the more important determinant of invasiveness although the underlying mechanisms are unclear.

Surprisingly, PXDN mRNA levels were also highly reduced in the KO (Figure 4) which suggests that the PXDN protein may play signalling roles and might be involved in feedback loops that impact gene transcription dynamics. Transcriptomics analysis of NZM40 PXDN KO vs WT cell lines indeed identified changes in a number of signalling pathways including WNT, PTK2, TGF-b signalling and the regulation of insulin-like growth factor transport and uptake by insulin-like growth factor binding protein, which are all pathways with implications in melanoma progression.

While PXDN is not a transcription factor, the vast changes in gene transcription following *PXDN* KO suggest that its expression is able to affect transcription factor activity. Interestingly, the transcriptional perturbations arising from *PXDN* KO encompassed a large number of genes encoding structural and functional ECM proteins, matrix metalloproteinases, as well as integrin and non-integrin membrane-ECM interacting proteins and cell junction proteins. These data highlight that PXDN is able to affect many aspects of ECM structure and function, regulation of cellular adhesion, invasion and degradation. The breadth of observed changes suggests that PXDN may affect several different transcription factors.

Additionally, our study provides convincing transcriptomic evidence suggesting that PXDN promotes an invasive phenotype. Phenotype plasticity, closely intertwined with non-mutational transcriptional and epigenetic reprogramming, is an emerging hallmark of cancer (42) and is referred to as phenotype switching in melanoma (2, 43). Next to epigenetic changes, cues from the tumour microenvironment (TME) like biochemical and biophysical properties are thought to play crucial roles in phenotype transition (44). PXDN has the potential to affect several properties of the TME, either via its catalytic activities (45, 46) or via protein-protein interactions, through its typical protein binding domains, which include a leucine-rich repeat domain, four C-like immunoglobulin domains and a von Willebrand factor type C module (47). Laminin, a major component of the basement membrane and TME, is the only so far identified binding partner of PXDN (15) and some laminins were downregulated in the *PXDN* KO. Specificity of oxidative modifications including collagen IV by HOBr generated by PXDN may be facilitated by transient protein-protein interactions. PXDN protein-protein interactions might also be involved in cell signalling processes.

Oxidants similar to HOBr have been shown to impact DNA methylation (48, 49), the activation of transcription factors (50) and signal transduction cascades (51, 52). One theory is that increased PXDN activity causes dysregulated oxidant generation, as is well documented in the cardiovascular context (53–59), where it affects cell signalling to promote oxidative stress, inflammation, ECM remodelling and endothelial dysfunction. PXDN activity has been shown to promote angiogenesis (60) and is upregulated in tissue fibrosis (16) which furthermore confirms its ability to shape the TME. Oxidative modifications of ECM components, including collagen IV cross-linking (13) may impact the structure and stiffness of the tumour ECM (19), promote proteolytic degradation (61), and affect cytokine and growth factor availability (62).

In conclusion, our study provides in depth transcriptomic and functional evidence that PXDN promotes the invasive melanoma phenotype. Transcriptomic changes following *PXDN* KO suggest that PXDN plays a prominent role in the reprogramming of the melanoma tumour ECM to promote invasion and tumour progression. Our data links PXDN to several so far unidentified cancer pathways and thereby sheds light on the function of PXDN in melanoma. Further studies will be required on additional cell lines, including additional *PXDN* knockouts and knock-ins, to elucidate detailed underlying mechanisms and studies in other cancer types will help to identify if these are common. Our results imply that PXDN may be an effective therapeutic target to decrease melanoma cell invasion. This could potentially be achieved with a specific PXDN inhibitor or therapeutic antibodies. To elucidate the best approach, it is important to understand whether PXDN acts via its catalytic activities, protein-protein interactions or a combination of both processes.

## Supporting information

Supplementary section

## Authors contributions

CCSD, bioinformatics analyses of transcriptomics and processed proteomics data, manuscript writing, editing, interpretation; AK, proteomics acquisition and data analyses; AD, CRISPR-Cas9 knockout design; PP, CRISPR-Cas9 knockout validation; KC, spheroid invasion assay guidance; NJM, bromotyrosine mass spectrometry work; SMH, bioinformatics guidance; MRC, interpretation, editing; CCW, conceptualization, funding acquisition, interpretation, editing; MPP, conceptualization, funding acquisition, cell line and knockout work, invasion assays, manuscript writing, editing, coordination, submission.

## Funding agencies

This research was supported (in part) by the Marsden Fund of the Royal Society of New Zealand (UOO1805), the Health Research Council of New Zealand, the Canterbury Medical Research Foundation, New Zealand; and the Cancer Research Trust New Zealand.

## Data availability statement

The authors confirm that the data supporting the findings of this study are available within the article and the supplementary section. All NZM40 WT and KO RNAseq data has been submitted to GEO (Gene Expression Omnibus).

## Abbreviations

PXDN: peroxidasin
EMT: epithelial-mesenchymal transition

