## Supplementary section for "Peroxidasin is associated with a mesenchymal-like transcriptional phenotype and promotes invasion in metastatic melanoma"

**Running title:** Peroxidasin promotes epithelial-mesenchymal transition in metastatic melanoma

**Key words**: peroxidasin, metastatic melanoma, cancer cell invasion, epithelial mesenchymal transition, phenotype plasticity, hypobromous acid, RNAseq, proteomics

**Abbreviations**: PXDN, peroxidasin; EMT, epithelial-mesenchymal transition;

**Contents:**

1. **Supplementary Materials & Methods:**

   CRISPR-Cas9 NZM40 *PXDN* knock-out generation

   LC-MS/MS Method for 3-bromotyrosine detection

   Label-free quantitative proteomics of NZM40 *PXDN* KO and WT

2. **Supplementary Figures:**

   Figure S1: CRISPR-Cas9 editing of NZM40 *PXDN* knockout

   Figure S2: Heatmap of Widmer Gene Sets of invasive and proliferative phenotype genes

   Figure S3: Overlapping Downregulated Genes in Low PXDN NZM Cells Lines and *PXDN* KO NZM40 vs WT

   Figure S4: Additional spheroid images

   Figure S6: Anti-PXDN antiserum validation

### 3. Supplementary Files

1.1 DEG Low vs High PXDN NZM.xlsx

1.2 Melanoma.Phenotypes.2.gmt.txt

1.3 Downregulated Pathways Reactome NZM.csv

1.4 Upregulated Pathways Reactome NZM.csv

4.1 DEG PXDN KO vs WT NZM40.xlsx

4.2 All Genes PXDN KO vs WT.xlsx

4.3 Downregulated Pathways Reactome NZM40.csv

4.4 Upregulated Pathways Reactome NZM40.csv

6.1 Lysate Proteomics.csv

6.2 Secretome Proteomics.csv

6.3 Downregulated Pathways Secretome.csv

6.4 Upregulated Pathways Secretome.csv

S3 Overlapping genes

**Supplementary Materials & Methods:**

**CRISPR-Cas9 NZM40 *PXDN* knock-out generation**

A knock-out cell line was generated by electroporating NZM40 with a plasmid containing Cas9 and a sgRNA targeting exon 10 of PXDN (Fig S1). The plasmid (pRP[CRISPR]-Neo-hCas9-U6>mPXDN[grna#22] VB201123-1155uqc) was designed and purchased from VectorBuilder. 200,000 cells were transfected with the Neon®Transfection system (1600V, 20 ms, 1 pulse) and seeded in antibiotic free medium after 10 min rest. 16 hours later 750 μM geneticin was added for four days to select for transfected cells. 100 units/mL of penicillin and 100 μg/mL of streptomycin were added on day three and geneticin was removed on day five. Cells were washed, tryspinized and 200 cells were seeded in a 15 cm plate. Single clone colonies were picked and expanded. DNA was extracted and 39 CRISPR-Cas9 edited clones were PCR amplified and sequenced (PXDN exon10 forward primer: 5'GTGAGGATGGGGCTGAGCCCTGGCTCTG3'; reverse primer: 5'AGAGGTGCTGGTTGGGGAGAGCGTCACGC3').

**LC-MS/MS Method for 3-bromotyrosine detection**

NZM40 WT and *PXDN* KO cells were grown in 6 well plates to full confluence. Medium was exchanged to Hank's Balanced Salt Solution and 500 μM bromide was added 10 min before 100 μM of hydrogen peroxide was added. Buffer was removed and cells were harvested in PBS after 60 min and sonicated. Cell lysates were analysed after digestion with pronase at 37 °C. Stable isotope dilution liquid chromatography tandem mass spectrometry (LC-MS/MS) was used for the detection and quantification of 3-bromotyrosine. Isotopically labelled standards ($^{13}C_6$-Tyr and $^{13}C_9$–3-Br-Tyr) were used to control for experimental variations and ionization. The two bromotyrosine isotopes (m/z 260 for the $^{79}$Br-containing isotope and m/z 262 for the $^{81}$Br-containing isotope) occurred at a 1:1 ratio and their peak areas were summed for quantification. Standard calibration curves using the ratios of unlabelled to labelled analytes were used for quantification. Standards and samples were analyzed using an on-line solid phase extraction method with a 6500 QTrap mass spectrometer coupled to an Infinity 1290 LC, utilizing a two-position six-port valve and a two-position ten-port valve. Standards and samples were stored on the autosampler tray at 5°C before being injected on to a Strata-X-CW 25 µm on-line extraction cartridge (20×2.0 mm) using 100% water containing 0.1% formic acid (Solvent A). The cartridge was then washed with 100% Solvent A for 1.5 min. Using 77% acetonitrile containing 0.1% formic acid (Solvent B) and 23% 100 mm ammonium formate (Solvent C), retained analytes were reverse eluted from the cartridge on to a pre-equilibrated Imtakt Intrada Amino Acid column (150×3.0 mm). Isocratic elution was carried out for 9 min, followed by a linear gradient to 100% Solvent C over 30 sec. The cartridge and column were then flushed with 100% Solvent C for 1.5 min, before being re-equilibrated with 77% Solvent B and 23% Solvent C for 3 min, followed by re-equilibration of the cartridge alone with 100% Solvent A for 1 min. The column oven temperature was set to 40°C and a flow rate of 0.25 mL $min^{-1}$ was used throughout. Samples were analyzed and the areas of bromotyrosine and tyrosine peaks were determined using Analyst 1.7.2. All species were quantified by fragmenting the singly-charged parent ion $[M+H]^+$, monitoring the fragment ion resulting from the loss of CH2O2 in positive-ion mode (Table S1), measuring the area under the curve of the resulting peak, and then relating to standard calibration curves.

**Table S1.** The m/z values for the singly-charged parent and fragment ions, and the optimised parameters that were used to quantify each analyte in LC-MS/MS experiments. DP (declustering potential), EP (entrance potential), CE (collision energy), CXP (cell exit potential).

| Analyte | Parent (m/z) | Fragment (m/z) | DP | EP | CE | CXP |
|---|---|---|---|---|---|---|
| Tyrosine ($H^+$) | 182.08 | 136.08 | 99 | 10 | 25 | 13 |
| Tyrosine ($^{13}C_6$) ($H^+$) | 188.10 | 142.10 | 99 | 10 | 25 | 13 |
| $^{79}$Bromotyrosine ($H^+$) | 259.99 | 213.99 | 66 | 10 | 25 | 13 |
| $^{81}$Bromotyrosine ($H^+$) | 261.99 | 215.98 | 66 | 10 | 25 | 13 |
| $^{79}$Bromotyrosine ($^{13}C_9$) ($H^+$) | 269.02 | 222.01 | 66 | 10 | 25 | 13 |
| $^{81}$Bromotyrosine ($^{13}C_9$) ($H^+$) | 271.02 | 224.01 | 66 | 10 | 25 | 13 |

**Label-free quantitative proteomics of NZM40 *PXDN* KO and WT**

Cell samples were lysed in 50 mM triethylammonium bicarbonate buffer, 5% sodium dodecyl sulfate using repeated sonication and vortex. Genomic DNA was degraded with a nuclease, Denarase (c-LEcta, Germany) in presence of 1 mM magnesium chloride. A BCA protein estimation assay (ThermoFisher Scientific, USA) was used to normalise the protein amount to 100 µg in all samples. Reduction and alkylation were carried out by 5 mM Tris(2-carboxyethyl)phosphine hydrochloride (TCEP) (Sigma-Aldrich, USA) and 10 mM iodoacetamide (IAM) (GE Healthcare, USA). Samples were then processed using an S-trap micro spin trap column (1) (ProtiFi, USA) according to the manufacturer's protocol (protifi.com/pages/protocols). The proteins on the column were tryptically digested, and cleaved peptides were eluted from the column for the proteomics analysis.

The volumes of cell medium samples containing the secretome were reduced using a 10 kDa MW cut-off Amicon spin-filter (Merck, USA). Before loading the samples to the filter unit, phenylmethylsulfonyl fluoride, a protease inhibitor, was added to the samples at a concentration of 1 mM. Once the sample volumes were reduced to approximately 100 µL, they were processed using a filter assisted sample processing (FASP) protocol (2). In brief, samples were on-filter reduced and alkylated with 5 mM TCEP and 10 mM IAM, respectively. Thereafter, a Bradford based protein estimation assay (BioRad, USA) was used to normalise the protein amount between samples to 25 µg. The samples were then tryptically digested, and the cleaved peptides were recovered from the filter unit.

Both cell and secretome samples were chromatographically separated on an in-house packed 20 cm emitter-tip column (75 µm ID fused silica tubing (CoAnn Technologies, USA) packed with 3 µM C-18 Luna material (Phenomenex, USA)) on an Ultimate 3000 uHPLC (Thermo Scientific, USA) coupled to the LTQ Orbitrap XL mass spectrometer (Thermo Scientific, USA). The peptides were eluted from the column with a reverse phase acetonitrile (ACN) gradient. Two different chromatography methods of 180 minutes and 120 minutes were used for the cell lysate and cell-media respectively due to the difference in protein content. The gradient for the cell pellet consisted of the following steps**:** 5 % to 25 % ACN in 133 minutes, 25 % to 40 % ACN in 15 minutes and 40 % to 99 % ACN in 7 minutes. Similarly, the gradient for the cell-media consisted of the following steps, 5 % to 25 % ACN in 81 minutes, 25 % to 40 % ACN in 11 minutes and 40 % to 99 % ACN in 4 minutes. In the mass spectrometer, peptides were measured at a resolution of 60000 @ m/z 400. The 11 strongest ms1 precursors between 400-2000 m/z were selected for collision induced dissociation (CID) fragmentation in the ion-trap. A normalised CID collision energy was set at 35 % with an AGC target of 2e5. Dynamic exclusion was enabled with 2 repeat counts during 90 seconds and an exclusion period of 180 seconds. Each biological repeat was measured in 3 technical replicates.

The resulting data from all samples were analysed with the Proteome Discoverer (PD) software (version: 2.5, Thermo Scientific, USA). The data was searched against the human proteome (downloaded: February 2022, Uniprot.org) with the Sequest HT search engine node inside the PD software. The search was set up to look for semi-tryptic peptides. In further search settings, carbamidomethyl cysteine was included as static modification and deamidation of asparagines and glutamines were included as variable modifications. The precursor mass tolerance and the maximum fragment mass error threshold was set at 10 ppm and 0.6 Da respectively. The FDR threshold was set at 1 % in the percolator node within the PD software and peptides and proteins within this threshold were classified as "high confidence". The resulting quantitative data was normalised on the sum of abundances from all peptides detected from all samples. The relative abundance of the proteins was calculated with the Top-3 approach (3) where the average abundance of the three most abundant peptides for a particular protein was used. The resulting abundance values were used to calculate the protein ratio between NZM40 *PXDN* KO to WT to generate a list of differentially abundant proteins. The data was exported to an Excel spreadsheet for further interpretation.

**Supplementary Figures:**

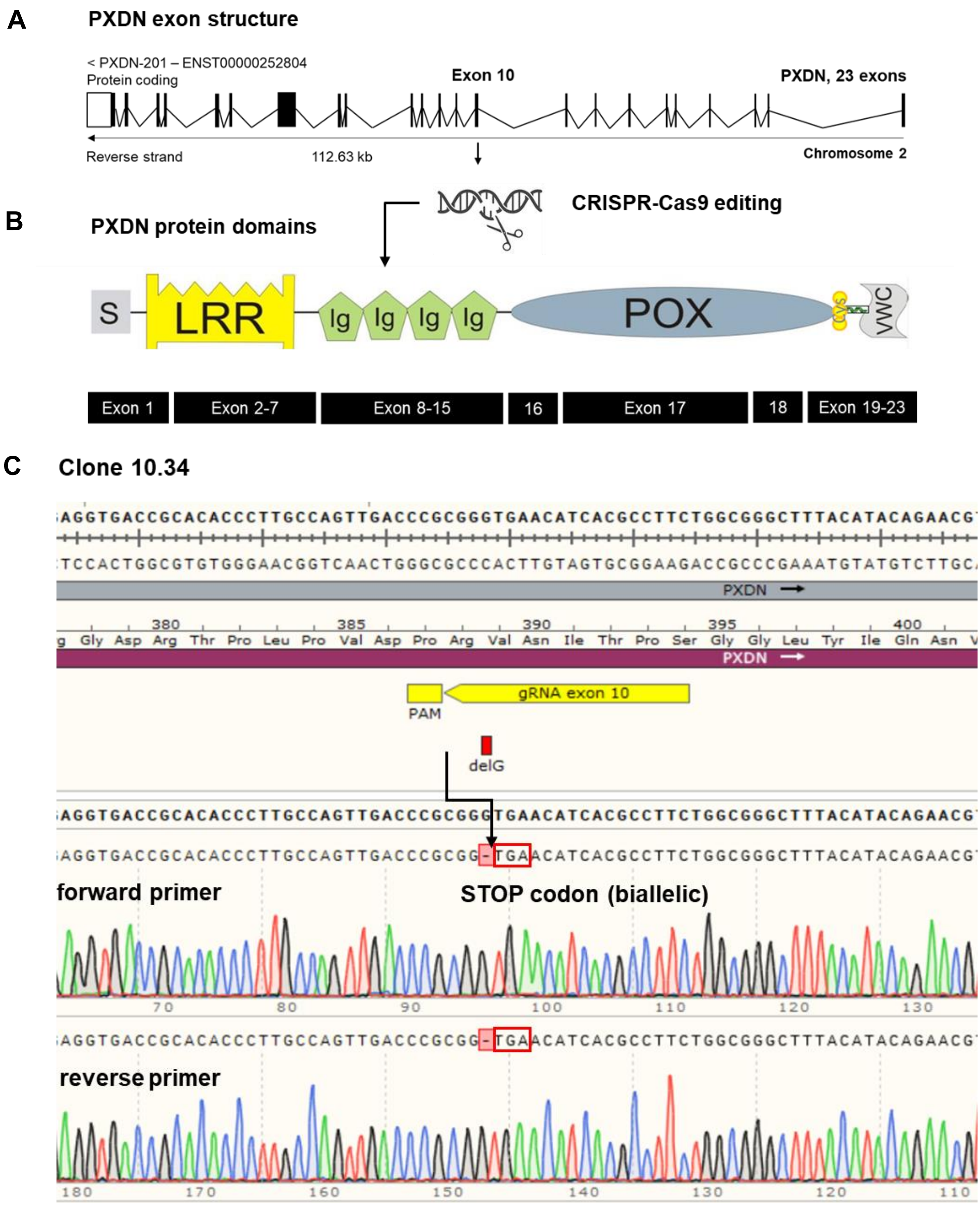


**Figure S1: CRISPR-Cas9 editing of NZM40 *PXDN* knockout. A:** Exon-intron-structure of *PXDN* gene located on the reverse strand of chromosome 2. gRNA was directed against exon 10 (arrow). **B:** PXDN protein domain structure depicting CRISPR-Cas9 editing site on the second Ig domain. **C:** PCR sequencing of clone 10.34 confirming the biallelic loss of one guanine nucleotide (highlighted in red) leading to a subsequent stop codon (TGA, red box).

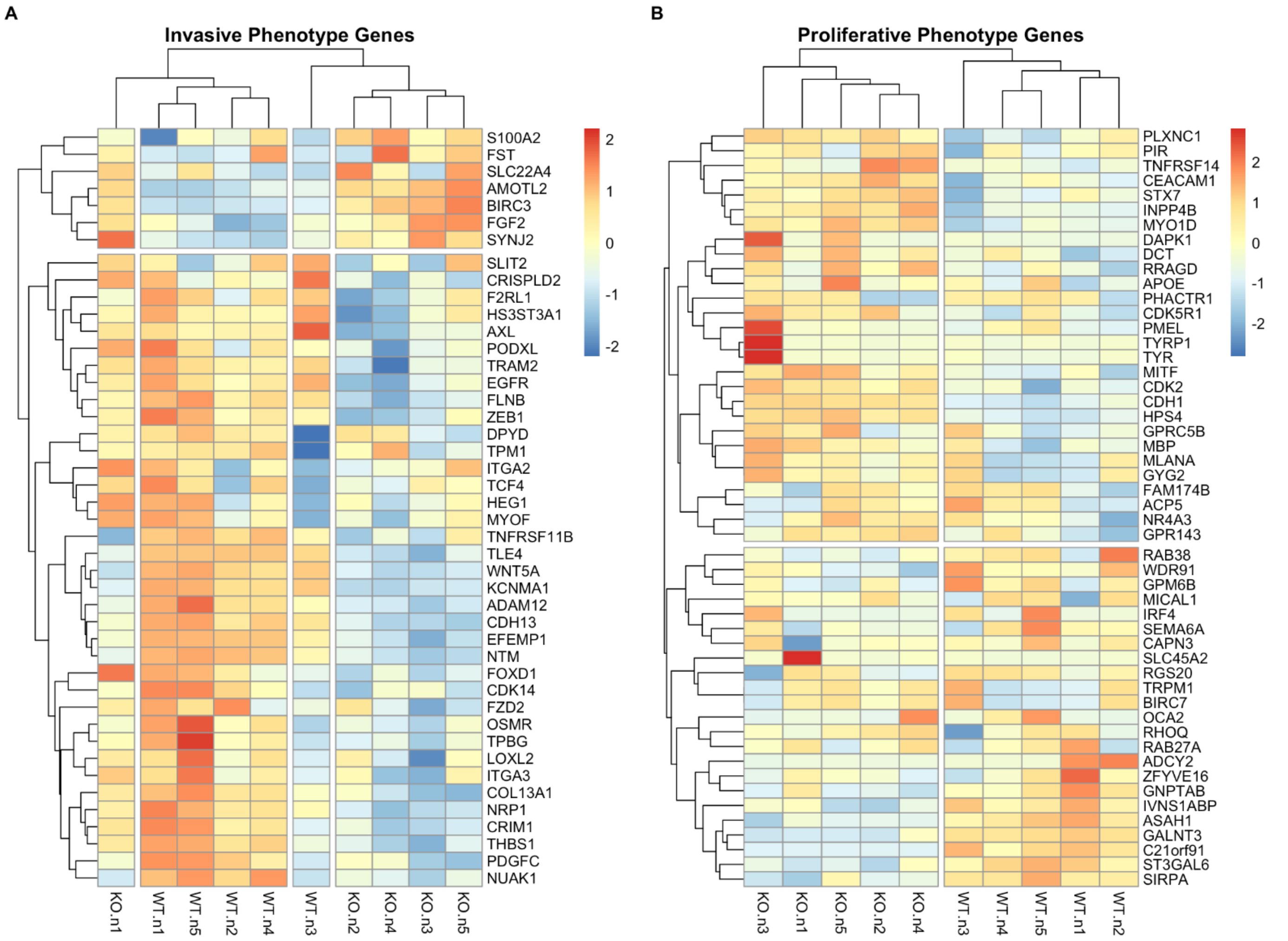


**Figure S2: Heatmap of Widmer Gene Sets of invasive and proliferative phenotype genes.** A and B: Heatmaps depicting the expression of genes identified as being linked to invasive or proliferative melanoma phenotypes based on Widmer et al. The majority of invasion related genes exhibit a decrease in expression in NZM cells following *PXDN* KO (A). In contrast, proliferative phenotype associated genes do not display such a compelling uniform change in expression (B).

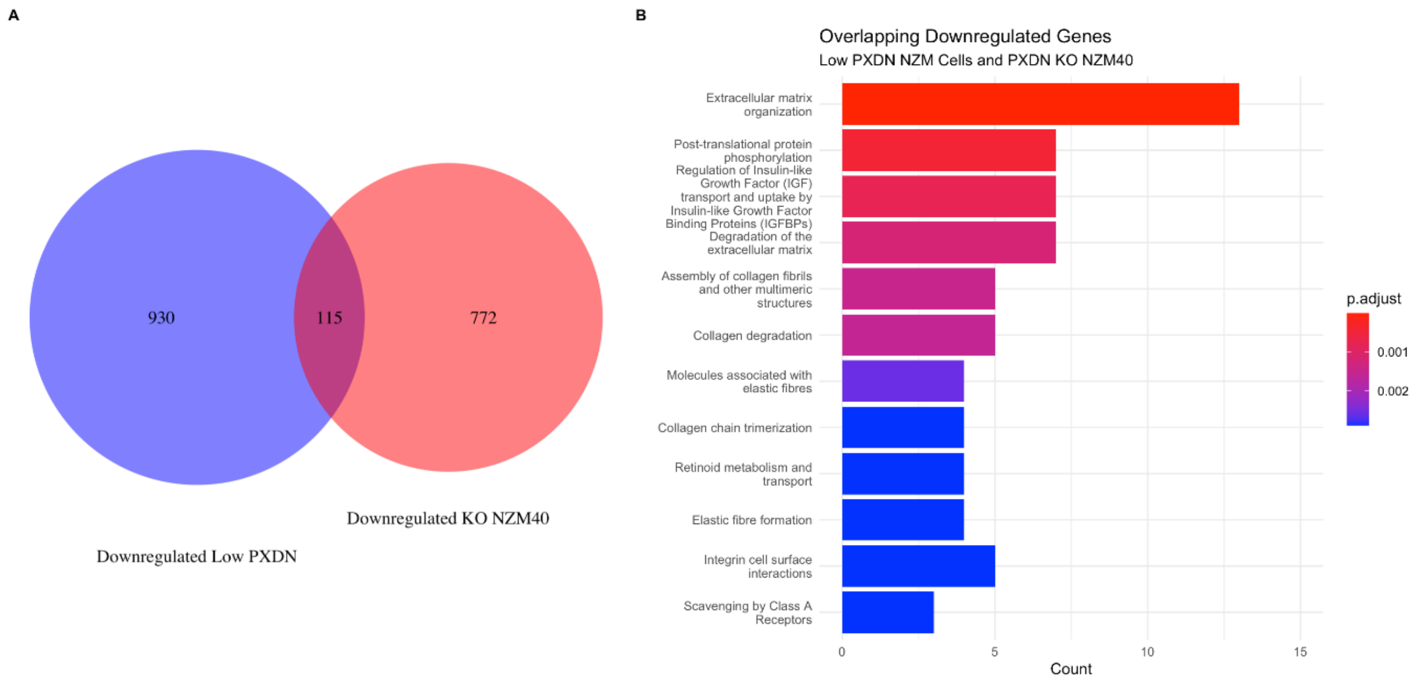


**Figure S3: Overlapping Downregulated Genes in Low PXDN NZM Cells Lines and *PXDN* KO NZM40 vs WT. A:** Genes were filtered to select downregulated genes (FDR <0.05, log2FC < 1) in either low vs high PXDN cell lines or NZM40 *PXDN* KO vs WT. 115 overlapped genes were identified and the list of genes is attached as a supplementary file. **B:** Performing overrepresentation analysis on these 115 genes identified pathways pertaining to the extracellular matrix and collagen assembly/degradation. A list of these genes is provided as a supplementary file along with a file of the overrepresentation analyses.

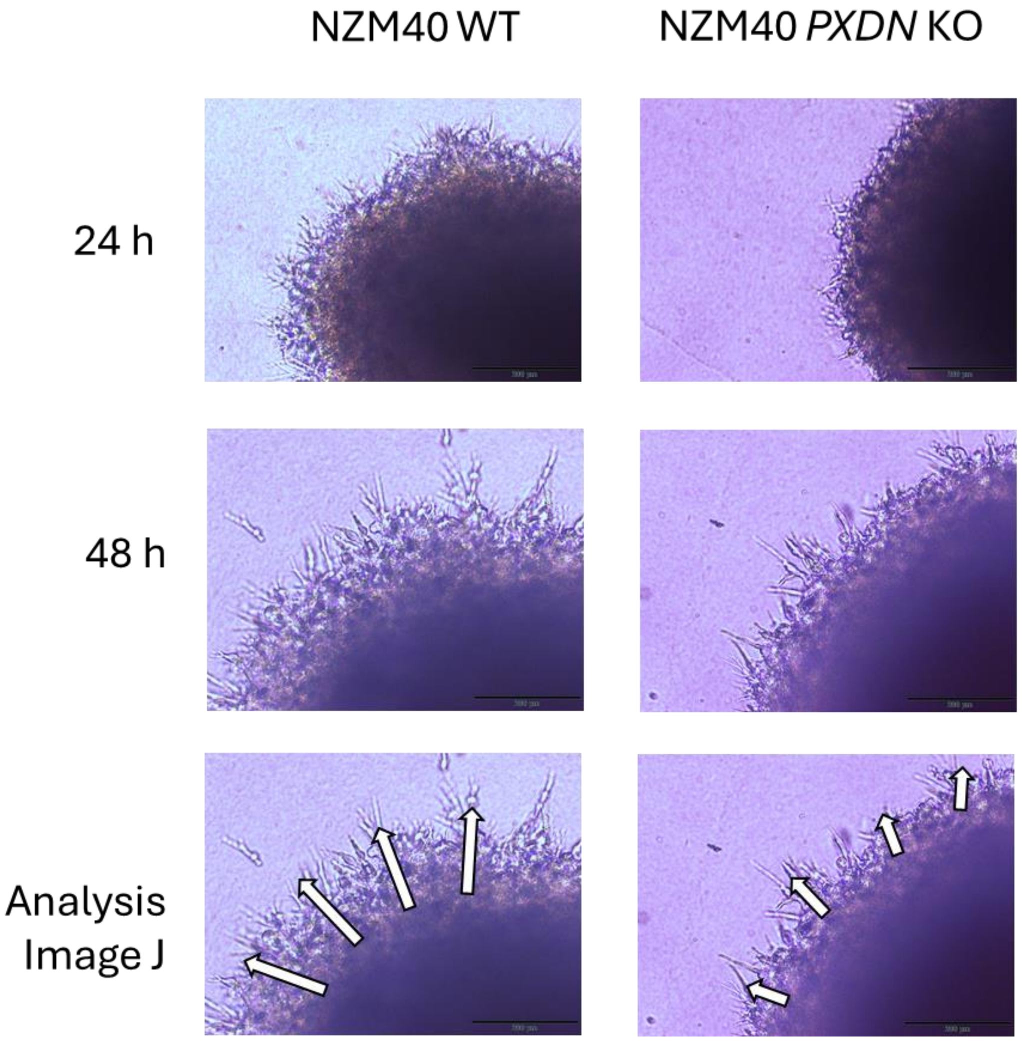


**Figure S4: Spheroids of NZM40 WT and *PXDN* KO.** Images taken at 24 and 48 hours and Image J analysis approach at 48-hour timepoint.

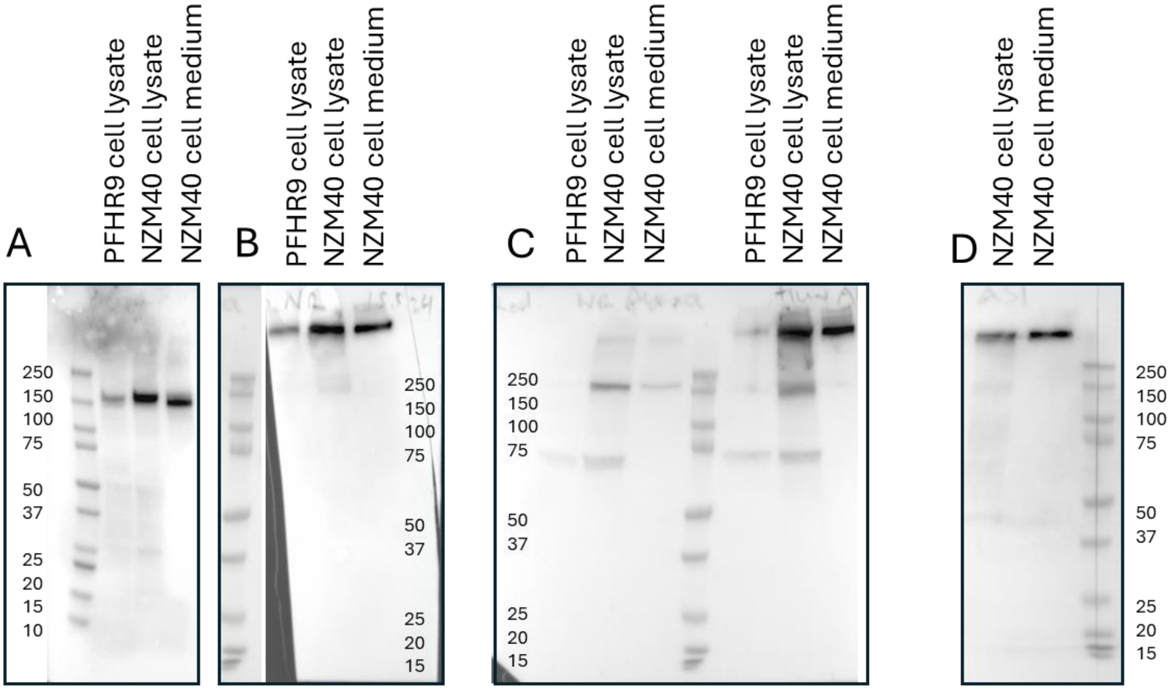


**Figure S5: Validation of anti-PXDN antiserum used in the ELISA (Figure 2A) and Abbexa anti-PXDN antibody used for western blotting in Figure 2B.** 50 ug of protein of PFHR9 (PXDN expressing mouse carcinoma cell line) and NZM40 cell lysates and 15 uL of NZM40 medium resolved on 4-15 % SDS-PAGE blotted onto PVDF membrane and probed with a mouse monoclonal anti-PXDN antibody kindly donated by Prof Miklos Geizst, developed in Budapest (4), under **A** reducing and **B** non-reducing condition. **C** Abbexa anti-PXDN antibody (abx101905) under reducing (left three lanes) and non-reducing condition (right three lanes). **D** In-house produced anti-PXDN antiserum under non-reducing conditions as used in the ELISA in Figure 2A. Under non-reducing conditions PXDN is resolved in its trimeric form of about 500 kDa (B, C right lanes, D), whereas under reducing conditions PXDN runs at its approximate monomeric molar mass of ~170 kDa. Depending on the antibody we also detect a monomeric and dimeric band at times under non-reducing conditions, which may represent the intracellular intermediates before full trimerization has occurred. PXDN is heavily glycosylated, which makes the protein appear at a slightly higher molar mass than expected and leads to decreased sharpness of the bands.

**Authors contributions:** CCSD, bioinformatics analyses of transcriptomics and proteomics data, manuscript writing, editing, interpretation; AK, proteomics acquisition and data analyses; AD, CRISPR-Cas9 knockout design; PP, CRISPR-Cas9 knockout validation; KC, spheroid invasion assay guidance; NJM, bromotyrosine mass spectrometry work; SMH, bioinformatics guidance; MRC, interpretation, editing; CCW, conceptualization, funding acquisition, interpretation, editing; MPP, conceptualization, funding acquisition, cell line and knockout work, invasion assays, manuscript writing, editing, coordination, submission.

**Conflict of interest:** The authors declare no conflict of interest.

**Funding agencies:** This research was supported (in part) by the Marsden Fund of the Royal Society of New Zealand (UOO1805), the Canterbury Medical Research Foundation, New Zealand; and the Cancer Research Trust New Zealand.

**Data availability statement:** The authors confirm that the data supporting the findings of this study are available within the article and the supplementary section. RNAseq data of NZM40 and NZM40 *PXDN* KO has been submitted to GEO (Gene Expression Omnibus).
